# Three years of insecticide resistance evolution and associated mechanisms in *Aedes aegypti* populations of Ouagadougou, Burkina Faso

**DOI:** 10.1101/2024.04.09.588738

**Authors:** Félix Yaméogo, Aboubacar Sombié, Manabu Oté, Erisha Saiki, Tatsuya Sakurai, Dimitri W. Wangrawa, Philip J. McCall, David Weetman, Hirotaka Kanuka, Athanase Badolo

**Affiliations:** Laboratoire d’Entomologie Fondamentale et Appliquée, Université Joseph Ki-Zerbo, Ouagadougou, Burkina Fas; Center for Medical Entomology, The Jikei University School of Medicine, Tokyo, Japan; Department of Tropical Medicine, The Jikei University School of Medicine, Tokyo, Japan; Laboratory Animal Facilities, The Jikei University School of Medicine, Tokyo, Japan; Université Norbert Zongo, Koudougou, Burkina Faso; Department of Vector Biology, Liverpool School of Tropical Medicine, Liverpool, UK

**Keywords:** *Aedes aegypti*, insecticide resistance, *kdr* mutation, pyrethroid, evolution, Burkina Faso

## Abstract

**Background:** Resistance to insecticides is spreading among populations of *Aedes aegypti*, the primary vector of important human arboviruses. The escalating insecticide resistance poses a significant threat to dengue vector control, with an expanding number of countries affected by the disease.

To gain a deeper insight into the evolution of insecticide resistance, it is essential to have longitudinal surveillance results, which are currently lacking, particularly from African *Ae. aegypti* populations.

Here we report on three-years of surveillance of *Ae. aegypti* susceptibility to insecticide resistance phenotypes and associated kdr mutations in Burkina Faso, a country with regular dengue outbreaks.

**Methods:** Ae. *aegypti* susceptibility to insecticides and the V410L, V1016I, and F1534C *kdr* target site mutations linked to pyrethroid insecticide resistance were monitored in Ouagadougou from 2016 to 2018. Larvae were collected from artificial containers at two sites and reared to adulthood in an insectary. Bioassays were conducted on female adults, alongside with a laboratory-susceptible strain, following standard WHO protocols. Allele-specific PCR genotyping assays were utilized to identify the V410L, V1016I, and F1534C *kdr* pyrethroid target site mutations.

**Results:** Bioassays revealed a high level of resistance to permethrin and deltamethrin that progressively increased over the three-year period in both localities. The 1534C mutation was nearly fixed throughout the three years at each locality, and while the closely related 410L and 1016I mutations did not vary between localities, their frequency notably increased from 2016 to 2018. Interestingly, *Ae. aegypti* populations in both areas remained susceptible to bendiocarb, fenitrothion, and malathion. Modeling the mortality data further confirmed the escalating resistance trend over the years and emphasized the significant role played by the three kdr mutations in conferring resistance to pyrethroids.

**Conclusion:** Mortality rates indicate that *Ae. aegypti* populations from Ouagadougou are becoming increasingly resistant to pyrethroid insecticides, likely due to an increase in the frequencies of the 410L and 1016I *kdr mutations*. Organophosphate insecticides are likely to be better alternative options for control.

## Introduction

Burkina Faso has a long history of dengue fever, with the first cases recorded in 1925 [1]. In 2015, several cases of dengue haemorrhagic fever were recorded in Ouagadougou [2], followed by a significant dengue outbreak in 2016, during which 1,256 suspected cases and 15 deaths were reported in the fourth quarter of the year [3]. In November 2017, a total of 9,029 cases and 18 deaths were reported throughout the country [4]

Dengue control and prevention relies primarily on vector control using larval source management [5] and chemical control of adult *Aedes* mosquitoes. This includes the use of organophosphates and pyrethroid insecticides for space spraying, and less commonly, pyrethroid-treated materials for personal or household protection [6]. Resistance to the four classes of insecticides most frequently used for vector control has been reported in *Aedes aegypti* populations worldwide, with particularly high prevalence in Latin American populations, showing broad-scale regional variation in at least some of the major resistance mechanisms involved [7].

*Aedes aegypti* has rarely been the target of focused vector control in Africa, except during periods of dengue outbreaks. Nonetheless, populations are often documented to exhibit resistance to pyrethroid insecticides, especially in west and central Africa [8, 9].

Pyrethroid resistance in Africa is typically associated with knockdown resistance (*kdr*) mutations in the voltage-gated sodium channel (Vgsc), especially 1534C, 1016I, and 410L [9, 10].

The F1534C variant is the most widespread mutation in *Ae. aegypti*, conferring resistance to deltamethrin and permethrin often in association with one or more of over 10 other mutations, such as V1016I and V410L [7, 11, 12].

The V1016I mutation confers resistance to pyrethroid insecticides only when associated with the F1534C mutation [11, 13], whereas V410L (also known as V419L) can confer resistance alone but is mostly found in linkage with V1016I [14, 15, 16]. The co-evolution of the V410L mutation with V1016I and F1534C can provide elevated levels of resistance to pyrethroids [16, 17]. After several years of resistance monitoring in Brazilian *Ae. aegypti*, [17] and [18] documented a geographical expansion and increase in the frequency of the 1534C and 1016I mutations, as well as evidence for increasing activity of the major classes of detoxifying enzymes.

Many reports on *Ae. aegypti* resistance to pyrethroid insecticides originate from Asia and the Americas, with data from Africa remaining sparse and longitudinal surveys lacking [19]. In this study, we report on the evolution of Ae. aegypti susceptibility to insecticides and the frequency of *kdr* mutations over three consecutive years in two localities during the same period of the year, following the 2016 dengue outbreak in Burkina Faso.

## Methods

### Study site, larval collection, and laboratory processing

Larvae and pupae of *Ae. aegypti* were collected from tires and drums during the rainy season for three consecutive years (2016-2018) from August to September. Two selected localities classified as peri-urban (Tabtenga: N12°22′05.7″ and W001°27′35.0″) and urban (1200 Logements N12°22′31.4″ and W001°29′51.3″), where dengue cases were previously reported, were chosen for mosquito sampling. The collected larvae and pupae were transferred to the insectary and reared to the adult stage using dried fish food (Tetramin®) until the adult stage. Emerged adult mosquitoes were fed a 10% sugar solution for 3-5 days, and females were used in bioassays. The insectary conditions were maintained at a temperature of 27±2°C, a relative humidity of 77±6%, and a 12:12 light: dark photoperiod.

### Adult susceptibility bioassays

Insecticidal bioassays were conducted following the WHO [20] tube test protocol. Five insecticides were tested: permethrin (0.75%), deltamethrin (0.05%), fenitrothion (1%), bendiocarb (0.1%), and malathion (5%). The discriminating doses of fenitrothion and bendiocarb correspond to those prescribed for *Ae. aegypti*, whereas those for permethrin, deltamethrin, and malathion recommended for *Anopheles gambiae*, and though very commonly used [7]], are higher than those for *Ae. aegypti*. Approximately 100 3–5-day-old unfed females were exposed to insecticide-impregnated papers (ordered from the Universiti Sains Malaysia WHO reference centre), and the number knocked down was counted every 10 minutes for one hour. Two tubes with non-impregnated paper, each containing 25 mosquitoes, were used as control. After the exposure period, mosquitoes were transferred to holding tubes for 24 hours, maintaining the same temperature and humidity conditions as mentioned earlier. The fully insecticide-susceptible Rockefeller strain of *Ae. aegypti* was used as a negative control to ensure the validity of the tests (100% mortality is expected for this strain). After 24 hours of observation, dead mosquitoes were counted, and the mortality rates were adjusted according to Abbott’s formula [21] when the control tube’s mortality rate ranged between 5 and 20%. Dead and alive mosquitoes were preserved in 1.5 ml micro-tubes over silica gel and stored at -20°C for molecular work.

Susceptibility tests were also performed following the CDC bottle protocol [22] to investigate potential metabolic resistance involvement in pyrethroid resistance through synergist effects, but interpretation was made according to WHO guidelines [20]. Permethrin and deltamethrin were used to coat bottles using acetone as a solvent for final concentrations of 0.015% (15µg) and 0.01% (10µg), respectively, as recommended for *Ae. aegypti*. For each bioassay using the synergist, approximately 150 non-blood-fed mosquitoes of 3-5 days were exposed first to the synergist (synergist exposure bottles) with 25 mosquitoes per bottle for 1 hour. After exposure to PBO, 100 mosquitoes were transferred to pyrethroid-coated bottles and 50 mosquitoes to acetone-coated bottles as insecticide control for 1 hour [22]. Two bottles were coated with acetone only as synergist controls. The insecticides and synergist PBO used were ordered from Sigma-Aldrich at technical grade purity (>98% purity). At the end of each exposure period, mosquitoes were removed and kept in cardboard cups for 24 hours of observation. At the end of the 24-hour observation time, dead and alive mosquitoes were preserved in 1.5 ml tubes over silica gel and stored at -20°C. Bioassay data from 2016 were published in [8] but are included for the purpose of comparison.

### DNA extraction and genotyping of V410L, V1016I and F1534C *kdr* mutations

DNA was extracted from mosquitoes using an ethanol precipitation method [23]. Genotyping of the F1534C kdr mutation was performed following the protocol of Li et al. [24] in a two-step PCR method with two common primers and one specific primer for each allele. Various genotypes of the 1016I mutation were detected using an allele specific PCR according to the protocols of Saavedra et al. [14] and Martins et al. [25], whereas genotyping of the V410L kdr mutation followed the protocol described by Granada et al. [13]. The wild-type and mutant alleles were identified in two separate PCR reactions.

For all PCRs conducted, each reaction tube contained 1 μl of target DNA, 6.25 μl of enzyme (AmpliTaq, Thermo Fisher Scientific), and 0.3 μM of each primer for a total volume adjusted to 12.5 μl with distilled water. Subsequently, 5μl of the PCR product was mixed with 6x loading buffer (New England BioLabs) and electrophoresed on a 1.5% TAE agarose gel for F1534C and V410L, and 3% TBE for V1016I. The gels were then stained with ethidium bromide and visualized under ultraviolet (UV) light.

### Statistical analysis

Mortality was calculated as the percentage of individuals that died within 24 h after 1-hour exposure to insecticide. Bioassay outcomes were interpreted according to WHO protocol [20]. Mosquitoes were considered resistant to an insecticide when mortality was less than 90%, and susceptible when mortality ranged between 98-100%. Resistance to insecticide is suspected when mortality is between 90-97% and requires subsequent testing for confirmation. The 95% confidence limits of mortality were calculated based on the method of [26]. A generalized linear model (glm) with a binomial linked function was fitted to the data to assess the effects of insecticide-locality-year and insecticide-locality, locality-year interactions on mosquito mortality. Moreover, the relationship between genotypes or haplotypes and phenotype (dead/alive) of mosquitoes was determined using a glm model using a binomial function. All analyses were performed using R software version 4.3.1. Significance level throughout was set at p<0.05.

## Results

### Adult susceptibility bioassay

The Rockefeller strain exhibited full susceptibility to all tested insecticides, leading to 100% mortality in WHO bioassays.

*Ae. aegypti* populations from 1200 Logements and Tabtenga remained fully susceptible to malathion and fenitrothion (100% mortality), during the three years of monitoring (Fig.1.a-b). The glm showed a significant difference (p<0.001) in mortality for fenitrothion in comparison with bendiocarb as reference, but not for malathion (p=0.98; Table 1).

**Table 1:** Parameters of Generalized linear model of *Ae. aegypti* mortality in 2016, 2017 and 2018 using WHO test. For each predictor, the incidence rate ratio, confidence limits and corresponding p-values are provided.

| Predictors | Odds Ratios | CI | P |
| --- | --- | --- | --- |
| (Intercept) | 16.63 | 11.42 – 24.90 | <0.001 |
| Insecticide [Bendiocarb] |  |  |  |
| Deltamethrin | 0.04 | 0.03 – 0.06 | <0.001 |
| Fenitrothion | 11.83 | 4.22 – 49.37 | <0.001 |
| Malathion | – | Inf – Inf | – |
| Permethrin | 0.02 | 0.01 – 0.03 | <0.001 |
| Locality [1200LG] |  |  |  |
| Tabtenga | 0.69 | 0.41 – 1.17 | 0.172 |
| Year [2016] |  |  |  |
| 2017 | 0.46 | 0.31 – 0.66 | **<0.001** |
| 2018 | 0.33 | 0.22 – 0.47 | **<0.001** |
| Insecticide [Bendiocarb] x Locality [1200LG] |  |  |  |
| Deltamethrin x Tabtenga | 0.78 | 0.44 – 1.39 | 0.394 |
| Fenitrothion x Tabtenga | 2.64 | 0.41 – 20.94 | 0.304 |
| Malathion x Tabtenga | – | – | – |
| Permethrin x Tabtenga | 0.86 | 0.49 – 1.53 | 0.604 |
| Locality [1200LG] x Year [2016] |  |  |  |
| Tabtenga x 2017 | 1.54 | 0.92 – 2.59 | 0.099 |
| Tabtenga x 2018 | 1.26 | 0.72 – 2.21 | 0.421 |

Significant predictors are highlighted in bold. Non-significant terms were interactions Insecticide\*Locality, Locality\*Year and Insecticide\*Locality \*Year. No survivors were recorded from the Malathion exposure.

Bendiocarb resistance was observed in both 1200LG and Tabtenga in 2016 (mortality<90%), but the status changed to suspected resistance in both localities in 2017 (mortality 90-97%), with a further reclassification in 2018 to susceptibility (mortality >98%; Figure 1.a-b).

**Figure 1:**
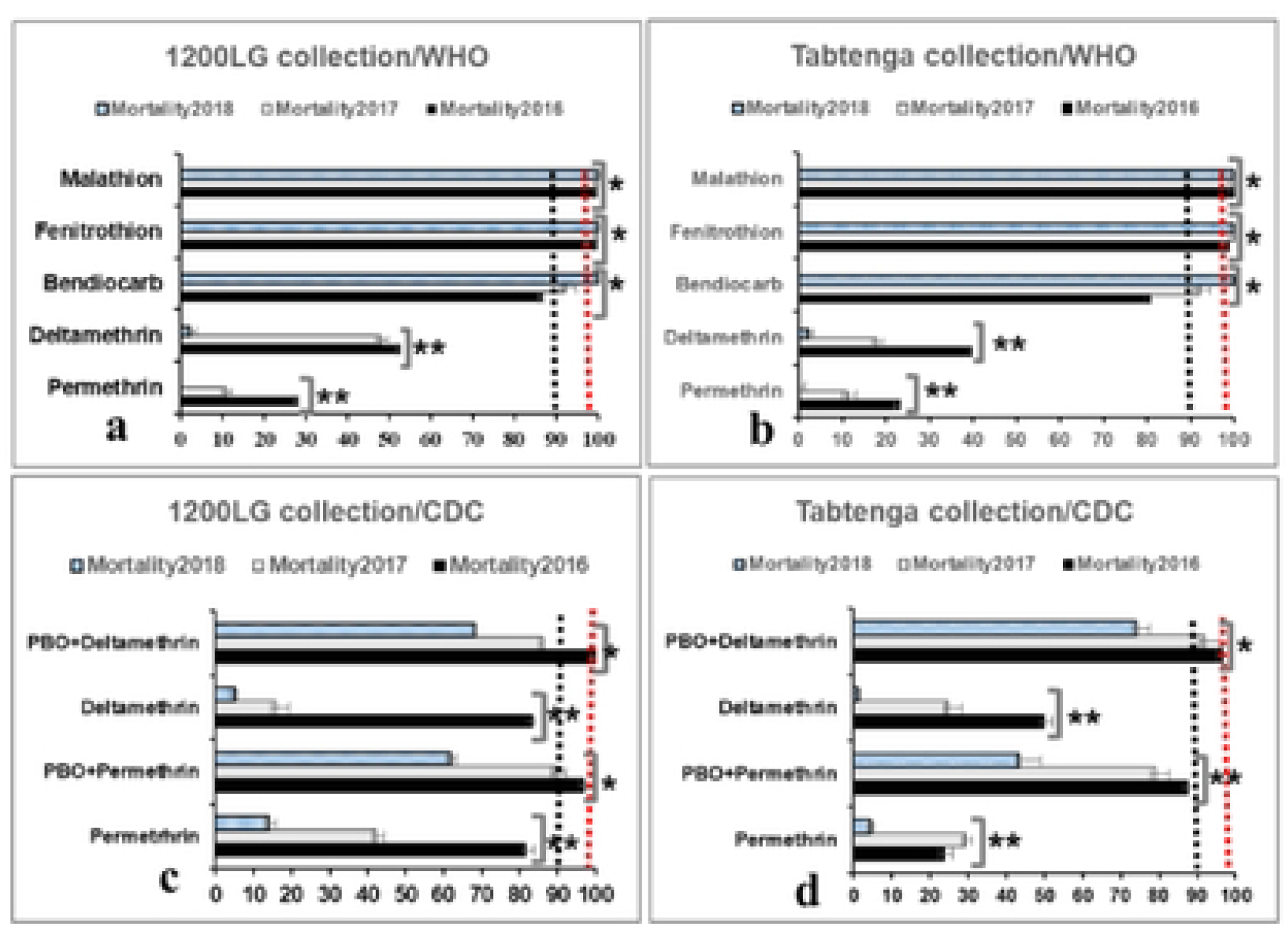
Mortality of *Ae. aegypti* adults exposed to insecticides. WHO tubes and CDC bottles 1 hour exposure to insecticides and synergist mortality (x-axis) and the 95% confidence limits of 1200LG (1200 Logements) and Tabtenga collection. The significance of the comparison between the three years for each locality is given: (**) significant, (*) Not significant). The red and black dashed lines indicate susceptibility and resistance thresholds, respectively.

WHO tubes bioassays revealed a high prevalence of resistance to permethrin and deltamethrin in each population across the three years with significantly decreasing mortality to both pyrethroids between 2016 and 2018 (p<0.001; Table 1 and Fig.1.a-b). Additionally, the GLM model showed a lesser effect of pyrethroid insecticide on mortality as indicated by odds ratios values compared to the reference bendiocarb (Table 1).

The trend of insecticide resistance using CDC bottles bioassays at recommended concentrations for *Ae. aegypti* mirrored WHO tube results for deltamethrin and permethrin, with generally decreasing mortalities over the three years for 1200 Logements and Tabtenga (Fig.1.c-d).

The combination of permethrin or deltamethrin with the PBO synergist partially restored the susceptibility. However, with increasing resistance, the degree of restoration also decreased across years. The glm revealed significant difference between mortalities in *Ae. aegypti* populations from 1200LG and Tabtenga during years 2016, 2017 and 2018, likely driven by slightly different responses to permethrin between sites across the years (Table 2).

**Table 2:** Parameters of generalized linear model of *Ae. aegypti* mortality in 2016, 2017 and 2018 using CDC bottles test. For each predictor, the incidence rate ratio, confidence limits and corresponding p-values are provided.

| *Predictors* | *Odds Ratios* | *CI* | *P* |
| --- | --- | --- | --- |
| **(Intercept)** | **3.92** | **2.83 – 3.53** | **<0.001** |
| Insecticide [Deltamethrin] |  |  |  |
| **PBO + Deltamethrin** | **29.17** | **17.34 – 50.65** | **<0.001** |
| **PBO + Permethrin** | **24.74** | **14.89 – 42.36** | **<0.001** |
| **Permethrin** | **2.16** | **1.39 – 3.41** | **<0.001** |
| Locality [1200LG] |  |  |  |
| **Tabtenga** | **0.20** | **0.13 – 0.31** | **<0.001** |
| Year [2016] |  |  |  |
| **2017** | **0.07** | **0.04 – 0.11** | **<0.001** |
| **2018** | **0.02** | **0.01 – 0.03** | **<0.001** |
| Insecticide [Deltamethrin] x Locality [1200LG] |  |  |  |
| PBO + Deltamethrin x Tabtenga | 1.44 | 0.71 – 2.90 | 0.315 |
| **PBO + Permethrin x Tabtenga** | **0.44** | **0.23 – 0.82** | **0.011** |
| **Permethrin x Tabtenga** | **0.29** | **0.16 - 0.51** | **<0.001** |
| Locality [1200LG] x Year [2016] |  |  |  |
| **Tabtenga x 2017** | **6.67** | **3.83 – 11.79** | **<0.001** |
| **Tabtenga x 2018** | **4.74** | **2.44 – 9.34** | **<0.001** |

Significant predictors are highlighted in bold. Non-significant terms were interactions Insecticide\*Locality\*Year.

### Evolution of F1534C, V1016I and V410L *kdr* mutations frequencies

The F1534C mutant exhibited high allelic frequency, nearing fixation in *Ae. aegypti* populations from both localities in 2016 and becoming fixed by 2018 (Figure 2.a-b). There were no wild-type homozygotes recorded in either locality, and heterozygotes were extremely uncommon (range 0 - 0.03).

**Figure 2:**
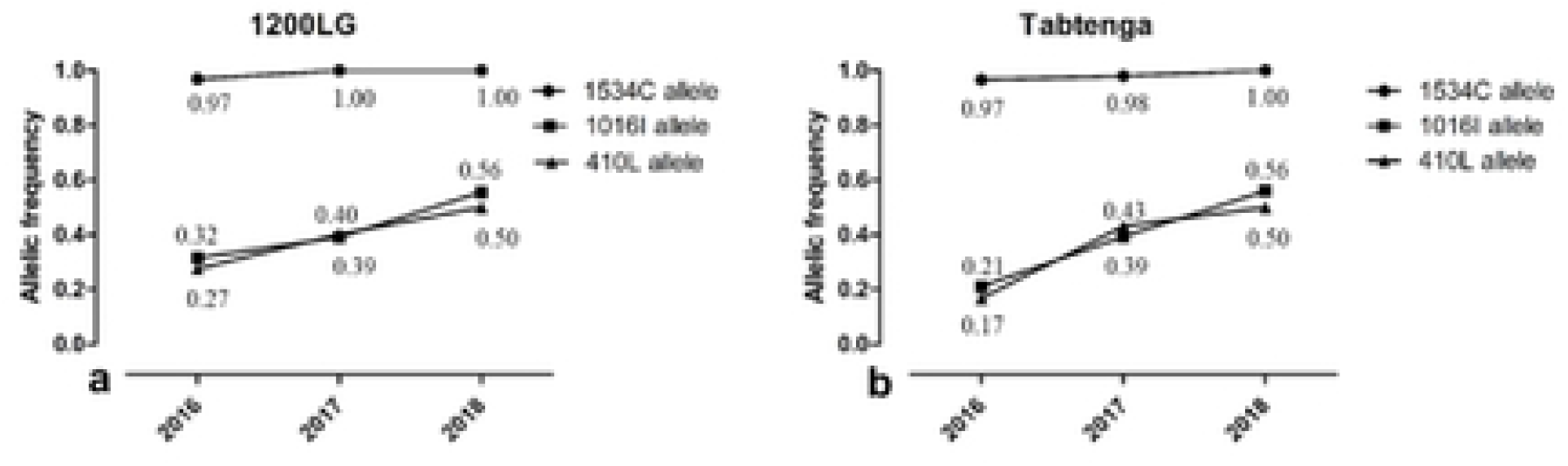
Evolution of *kdr* mutant allele frequencies (1534C, 1016I and 410L) in *Ae. aegypti* collected in 1200LG and Tabtenga during 2016, 2017 and 2018.

The frequency of the 1016I allele increased between 2016 and 2018 in both Tabtenga and 1200LG. The genotypic frequency of homozygous 1016I/I was lower in *Ae. aegypti* from 1200LG and Tabtenga in 2016, with the frequency of this genotype increasing in 2017 and 2018 in both localities (SFigure 1b). The 410L allele frequency also increased from 2016 to 2018 in 1200LG (Figure 2.a) and in Tabtenga during the same period (Figure 2.b). The genotype frequency of homozygous 410L/L increased between 2016 and 2018 for populations of 1200LG (SFigure 1.a) and Tabtenga (SFigure 1.d). The heterozygote (410V/L) frequency also increased in both localities from 2016 to 2018, in association with the wild allele (V410) frequency decreasing from 2016 to 2018 (SFigure 1.a-d).

### Evolution of F1534C, V1016I and V410L haplotypes, genotypes and their association with pyrethroid resistance

Five out of a total of eight possible haplotypes (CIL, CIV, CVL, CVV and FVV) were recorded from the genotypes of the three variants in the two localities, during 2017 and 2018 in *Ae. aegypti* exposed to permethrin and deltamethrin (Table 3). Data from 2016 are published [8], though without detection of the V410L *kdr* mutation at that time.

**Table 3:** Glm of survivor haplotype number of *Ae. aegypti* to permethrin and deltamethrin from 1200LG and Tabtenga collected in 2017 and 2018. For each predictor, the incidence rate ratio, confidence limits and corresponding p-values are provided.

| *Predictors* | *Odds Ratios* | *CI* | *P* |
| --- | --- | --- | --- |
| **(Intercept)** | **0.50** | **0.26 – 0.93** | **0.032** |
| Insecticide [Delta] |  |  |  |
| **perm** | **14.45** | **4.89 – 54.12** | **<0.001** |
| Locality [1200LG] |  |  |  |
| Tab | 0.54 | 0.24 – 1.23 | 0.146 |
| Year [2017] |  |  |  |
| **2018** | **22.87** | **6.06 – 151.09** | **<0.001** |
| Haplotypes [CVV] |  |  |  |
| **CIL** | **2.88** | **1.52 – 5.61** | **0.001** |
| CIV | 1.57 | 0.26 – 13.39 | 0.642 |
| FVV | 3.69 | 0.14 – 98.33 | 0.370 |
| Insecticide [delta] × Locality [1200LG] |  |  |  |
| perm × Tab | 0.32 | 0.07 – 1.28 | 0.121 |
| Locality [1200LG] × Year [2017] |  |  |  |
| Tab × 2018 | 2.76 | 0.29 – 26.58 | 0.350 |

The triple mutation 1534C/1016I/410L haplotype (CIL) and the single (1534) mutant haplotype CVV were the most prevalent in *Ae. aegypti* populations of the two localities (S1table), while CVL was rare. Only the CIL haplotype increased the survival of the mosquito (approximately threefold) compared to the reference CVV. The other haplotypes, CIV and FVV did not show a significant difference in survival to both permethrin and deltamethrin (p>0.05) (Table 3). Sample size limited the assessment of contribution of most haplotypes’ associations to insecticide resistance (Table 3).

The frequency of the CIL/CIL triple mutant genotype increased from 2016 to 2018 (S1table) in 1200LG and Tabtenga, whereas the CVV/CVV genotype frequency decreased in both localities between 2016 and 2018 (S1table). The wild type genotype FVV/FVV was absent and the FVV/CVV was infrequent during the first two years, and absent in 2018. As the 1534C allele is almost fixed, a glm of the genotypes based on the two mutations 1016I and 410L showed that the IL/IL genotype increased 6 times the survival of Aedes mosquitoes compared to the reference genotype VV/VV (S2Table).

## Discussion

Pyrethroid resistance in *Ae. aegypti*, reported in 2016 and 2017, was linked to a high frequency of the 1016I and 1534C kdr mutations [8, 24]. Our current study presents a three-year follow-up, showing an increasing insecticide resistance to pyrethroid insecticides associated with the 1534C, 1016I, and 410L kdr mutations in *Ae. aegypti* populations from Tabtenga and 1200LG in Ouagadougou. These populations exhibited high and increasing pyrethroid resistance over the surveillance period, with incomplete restoration of susceptibility even when the PBO synergist is associated with pyrethroids. The first documented West African case of *Ae. aegypti* pyrethroid resistance was reported in Ghana, involving the F1534C kdr mutation and the V1016I mutations [27]. Subsequently, resistance was also observed in Cameroon [28], albeit without the kdr mutation. Recent studies in Ghana and Cameroon have reported significant pyrethroid resistance linked to kdr mutations [31, 32]. Because previous studies in these countries were not carried out in the same localities, the spatial variation may be a confounding factor in the temporal increase in insecticide resistance and kdr mutation allelic frequency recorded. The current study has the advantage of being carried out in the same localities and at the same period over three consecutive years. That allows for a consistent demonstration of the escalating resistance of *Aedes aegypti* to pyrethroid insecticides and the associated kdr mutations over time in the two localities. The synergist PBO partially restored susceptibility to deltamethrin and permethrin in *Ae. aegypti*, indicating the involvement of cytochrome P450 monoxygenase activity in pyrethroid resistance, as observed in mosquitoes from the same localities with overexpression of P450 candidate genes [8]. The activity of PBO also decreased over the three years, correlating with declining permethrin and deltamethrin mortality when PBO was associated, suggesting a combination of target site and metabolic resistance.

Resistance was maintained for the three years of surveillance, and even more, the survivorship of *Ae. aegypti* to these doses increased. The distribution of the V410L, V1016I, and F1534C genotypes has shifted over the years, showing a rise in the allelic frequency of 410L and 1016I, while the 1534C mutation has nearly reached fixation in the population. Previous studies have established a link between insecticide application and the dynamics of resistance in *Ae. aegypti* [18], particularly the development of resistance due to prolonged insecticide use in dengue vector control campaigns [18, 33].

The haplotype 1534C/1016I/410L is significantly associated with permethrin and deltamethrin survivors at both the 1200LG and Tabtenga collections, indicating that the presence of this triple mutation induces pyrethroid resistance in *Ae. aegypti*. Similar findings were highlighted in Mexico, where the association between the triple mutation and pyrethroid resistance was observed [14]. Another study conducted in Mexico from 2000 to 2012 [32], reported an increase in the frequencies of the 1534C and 1016I mutations in *Ae*. aegypti, reflecting a similar trend observed in the current study. The 410L mutation, either alone or in combination with 1534C, confers high levels of resistance [14], making its emergence a challenge for vector control and compromising the use of pyrethroids [16]. The evolution of this triple mutation over a three-year period in the current study could largely explain the intensification of resistance observed with deltamethrin and permethrin.

The status of *Ae. aegypti* populations’ resistance or susceptibility to bendiocarb is not entirely clear in the current study. Resistance decreased progressively from 2016 (Mortality < 90%) to full susceptibility in 2018 in the two localities. However, an intensity test of 2 hours exposure to the diagnostic dose of bendiocarb did not confirm the 2016 resistance results in Tabtenga and 1200LG [8]. Additionally, Sombié et al. [23] confirmed the susceptibility to carbamates in Somgandé, while in 2017, resistance was recorded in the same locality, indicating spatial heterogeneity in carbamate resistance.

*Aedes. aegypti* has not been targeted by vector control measures in Burkina until the 2016 outbreak where malathion was used in spatial and indoor residual spray in dengue hotspots. There has been a shift from susceptibility to full resistance within a three year period for *Anopheles gambiae* in an area without any other change than the free distribution of insecticide treated bednets [33]. Several studies have made the link between bednets use and development of resistance in malaria vectors [36, 37] and in *Culex quinquefasciatus* [36]. LLINs were distributed in 2010, 2013, 2014 and 2016 in Burkina Faso increasing bednets coverage to 85% of the population [37]. Considering socio-economic conditions, it is likely that Tabtenga and 1200LG differ in the use of insecticide coils and sprays. Urban agricultural with use of insecticides in gardening may constitute additional selection pressure for *Ae. aegypti* resistance to insecticides [38]. However, the absence of organophosphate resistance while using these insecticides in Ouagadougou in agriculture and sprays [8, 10], supports the idea of resistance selection by insecticide impregnated bednets. Additionally, insecticide used in agriculture may have a limited effect as resistance pressure on *Ae. aegypti*, given that it primarily breeds in artificial containers in urban settings, even if physicochemical factors that affect mosquito body size [39] may also play a role in selecting resistance to insecticides [40]. Recent studies have shown that *Ae. aegypti* exhibits exophilic but more likely endophagic behaviour [41] meaning that insecticide-treated bednets and other indoor-used insecticides may have also contributed to the development of insecticide resistance.

## Conclusion

*Ae. aegypti* maintained resistance to pyrethroid insecticides over the three years of surveillance, with resistance levels increasing over time. The F1534C, V410L, and V1016I *kdr* frequencies increased, with the F1534C *kdr* mutation reaching fixation, and the homozygotes for both V410L and V1016I surviving pyrethroid insecticide exposure. Ae. aegypti remained susceptible to fenitrothion and malathion at the tested doses. These findings have significant implications for vector control planning against arbovirus vectors in Burkina Faso and may provide valuable insights for other African countries with limited data on insecticide resistance. Given the escalating insecticide resistance, urgent attention is required to develop insecticide resistance (IR)-mitigating strategies to prevent arbovirus outbreaks.

## Availability of data and materials

All data generated and/or analyzed during this study are included in this published article. All authors have read and approved the final manuscript.

### Ethics approval and consent to participate were obtained for this study. All necessary ethical guidelines and protocols were followed to ensure the protection of human subjects involved in the research

Not applicable.

## Consent for publication

Not applicable

## Author’s details

^1^Laboratoire d’Entomologie Fondamentale et Appliquée, Département de Biologie et Ecologie Animales, Université Joseph Ki ZERBO, Ouagadougou, Burkina Faso.

## Authors’ contributions

Field collections: FY, AS, AB. Laboratory experiments: FY, AS, AB. Reagents and laboratory materials:…, Statistical analysis: FY & AB.

**S1 Table:** Co-occurrence of F1534C, V1016I and V410L mutations in *Ae. aegypti* populations. The number of each genotype is provided, along with its frequency in parentheses. The probability of comparing the genotypes distribution over the three years for each locality was determined using the Fisher Exact test.

| Genotypes | 1200LG |  |  | Tabtenga |  |  |
| --- | --- | --- | --- | --- | --- | --- |
|  | 2016 | 2017 | 2018 | 2016 | 2017 | 2018 |
| CIL/CIL | 1 (0.020) | 8 (0.22) | 9 (0.20) | 2 (0.04) | 10 (0.19) | 12 (0.24) |
| CIV/CIL | 0 (0.00) | 0 (0.00) | 1 (0.02) | 0 (0.00) | 1 (0.02) | 2(0.08) |
| CIV/CIV | 0 (0.00) | 0 (0.00) | 0 (0.00) | 0 (0.00) | 1 (0.02) | 1 (0.08) |
| CVL/CIL | 0 (0.00) | 0 (0.00) | 0 (0.00) | 0 (0.00) | 0 (0.00) | 2 (0.04) |
| CVV/CIL | 23 (0.47) | 10 (0.28) | 23 (0.52) | 10 (0.23) | 23 (0.44) | 20 (0.40) |
| CVV/CIV | 4 (0.08) | 0 (0.0) | 6 (0.14) | 4 (0.09) | 1 (0.02) | 1 (0.02) |
| CVV/CVL | 0 (0.00) | 1 (0.03) | 0 (0.00) | 0 (0.00) | 0 (0.00) | 0 (0.00) |
| CVV/CVV | 18 (0.37) | 17 (0.47) | 5 (0.11) | 26 (0.58) | 14 (0.27) | 9 (0.18) |
| FVV/CIL | 2 (0.04) | 0 (0.00) | 0 (0.00) | 1 (0.02) | 1 (0.02) | 0 (0.00) |
| FVV/CVV | 1 (0.02) | 0 (0.00) | 0 (0.00) | 2 (0.04) | 1 (0.02) | 0 (0.00) |
| Fisher exact test | P<<0.001 |  |  | P=0.001 |  |  |

**S2 Table:** Glm of *survivor* genotypes number *of Ae. aegypti* to permethrin and deltamethrin from 1200LG and Tabtenga collected in 2017 and 2018. For each predictor, the incidence rate ratio, confidence limits and corresponding p-values are provided.

| **Predictors** | **Odds Ratios** | **CI** | **P** |
| --- | --- | --- | --- |
| (Intercept) | 0.45 | 0.15 – 1.25 | 0.138 |
| Insecticide [delta] |  |  |  |
| **perm** | **17.20** | **3.66 – 133.92** | **0.001** |
| Locality [1200LG] |  |  |  |
| Tab | 0.50 | 0.14 – 1.65 | 0.259 |
| Year [2017] |  |  |  |
| **2018** | **39.11** | **5.19 – 943.55** | **0.003** |
| Genotypes [VV/VV] |  |  |  |
| **IL/IL** | **6.47** | **1.62 – 30.66** | **0.012** |
| IV/IL | – | – | 0.996 |
| IV/IV | – | – | – |
| VL/IL | – | – | 0.998 |
| VV/IL | 1.50 | 0.50 – 4.55 | 0.467 |
| VV/IV | 0.21 | 0.02 – 2.43 | 0.191 |
| VV/VL | – | – | – |
| Insecticide [delta] × Locality [1200LG] |  |  |  |
| perm × Tab | 0.31 | 0.03 – 2.32 | 0.275 |
| Locality [1200LG] × Year [2017] |  |  |  |
| Tab] × 2018 | 2.18 | 0.07 – 69.76 | 0.622 |

**S1 Figure.**
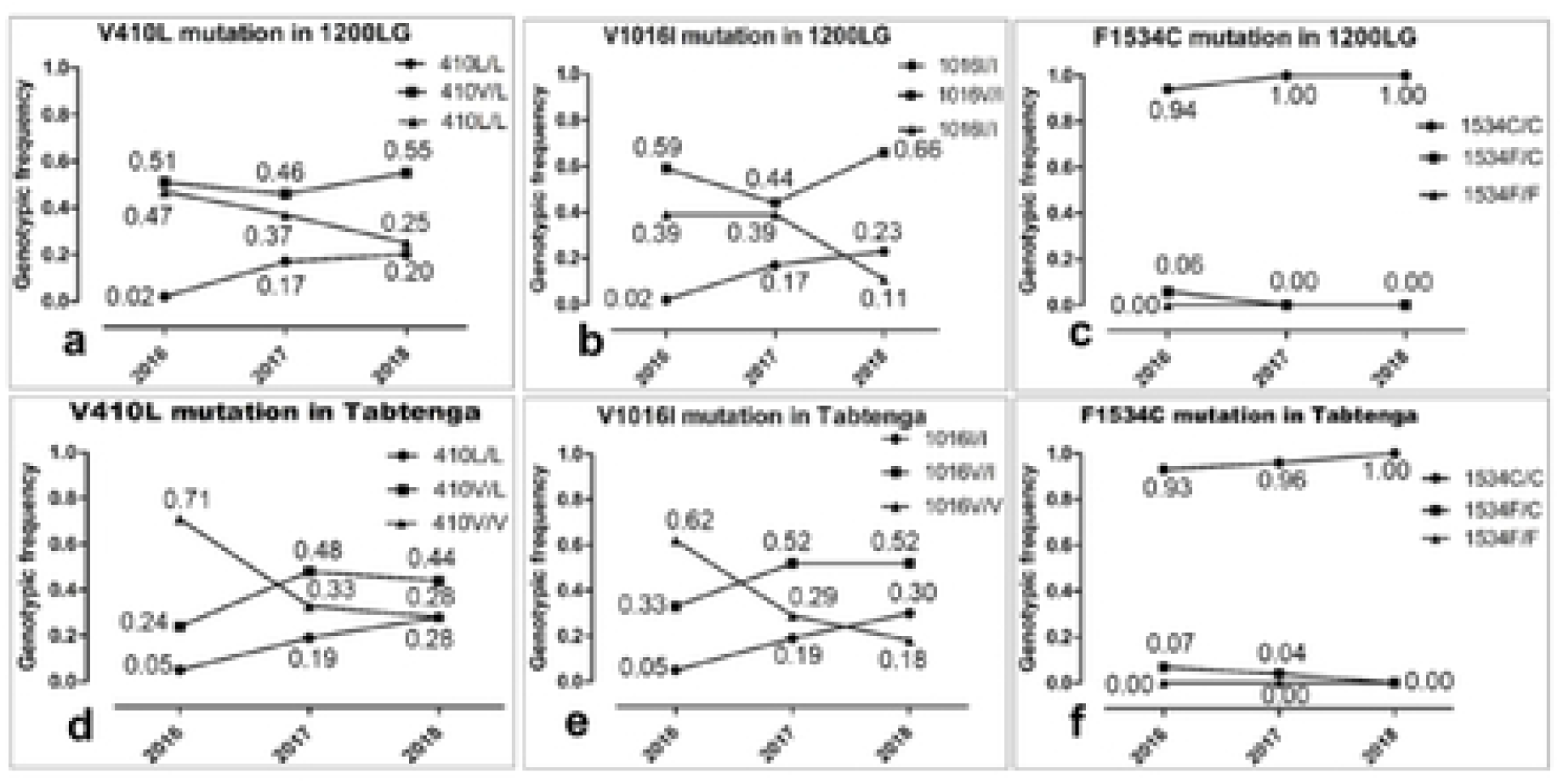
Evolution of genotypic frequencies of F1534C, V1016I and V410L mutations in *Ae. Aegypti* mosquitoes from 1200LG and Tabtenga during 2016, 2017 and 2018.

